## Supplemental figures for "The genetic and dietary landscape of the muscle insulin signalling network"

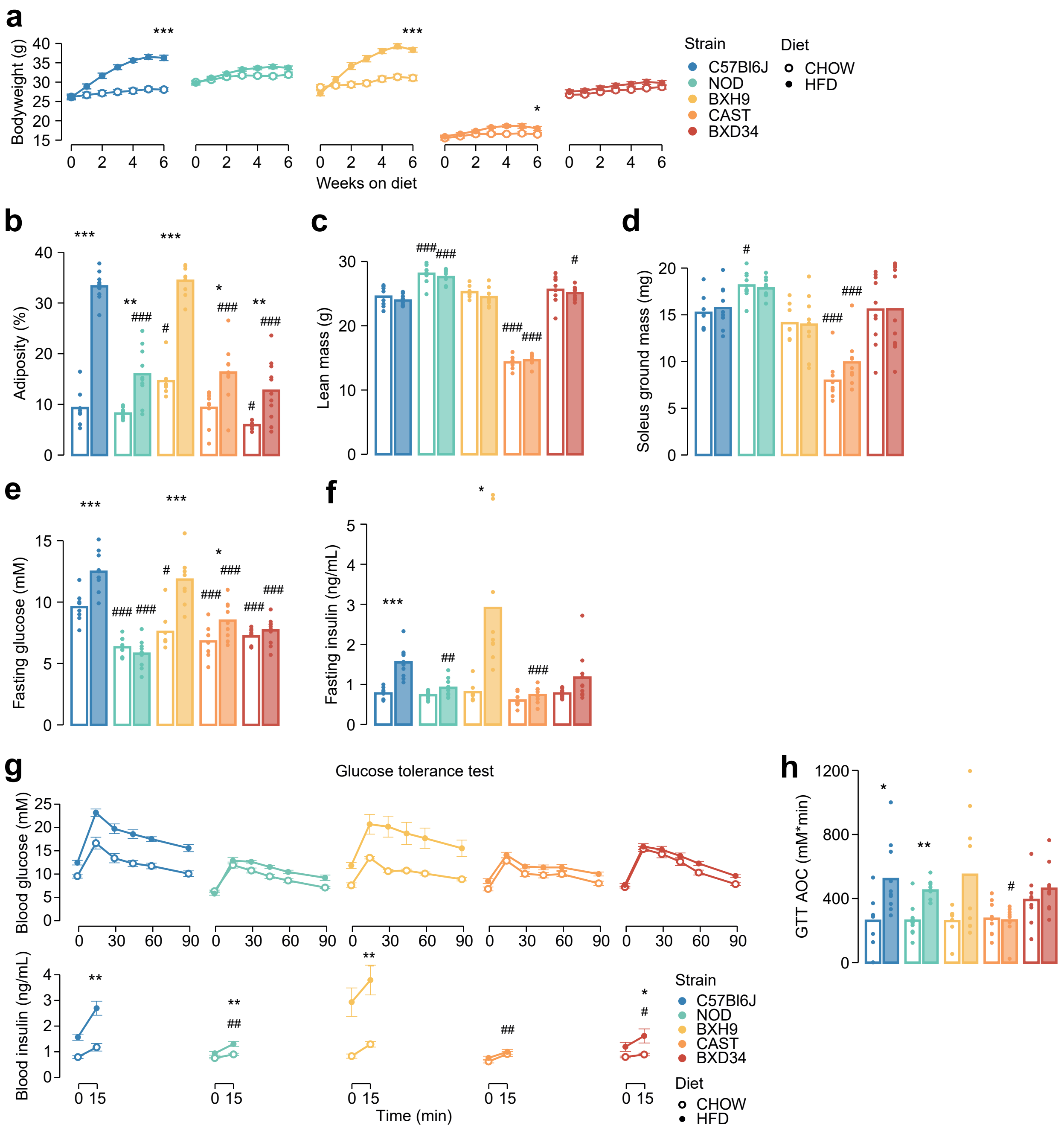


**Figure S1: Genetics and diet alter morphometric and metabolic phenotypes.** Related to Fig. 1. **a)** Mouse bodyweight was measured during a six-week diet regimen. Two-sided t-tests were performed to compare HFD to CHOW within each strain after six weeks, following Benjamini-Hochberg p-value adjustment (*). **b-d)** Measurement of **b)** adiposity, **c)** lean mass, **d)** ground soleus mass, **e)** fasting blood glucose, and **f)** fasting blood insulin at the end of the diet regimen. **g)** At the end of the diet regimen a glucose tolerance test was performed. **h)** The area of the blood glucose curve (GTT AOC) was calculated. In **b-h)**, two-sided t-tests were performed to compare HFD to CHOW within each strain (*) or to compare each strain to C57Bl6J within either diet (#). P-values were adjusted by the Benjamini-Hochberg procedure. Error bars indicate SEM. In **g)** t-tests were only performed on 15-minute blood insulin levels. No comparisons across strains on CHOW were significant. n = 8-11 biological replicates. */#: $0.01 \leq p < 0.05$, **/##: $0.001 \leq p < 0.01$, ***/###: $p < 0.001$

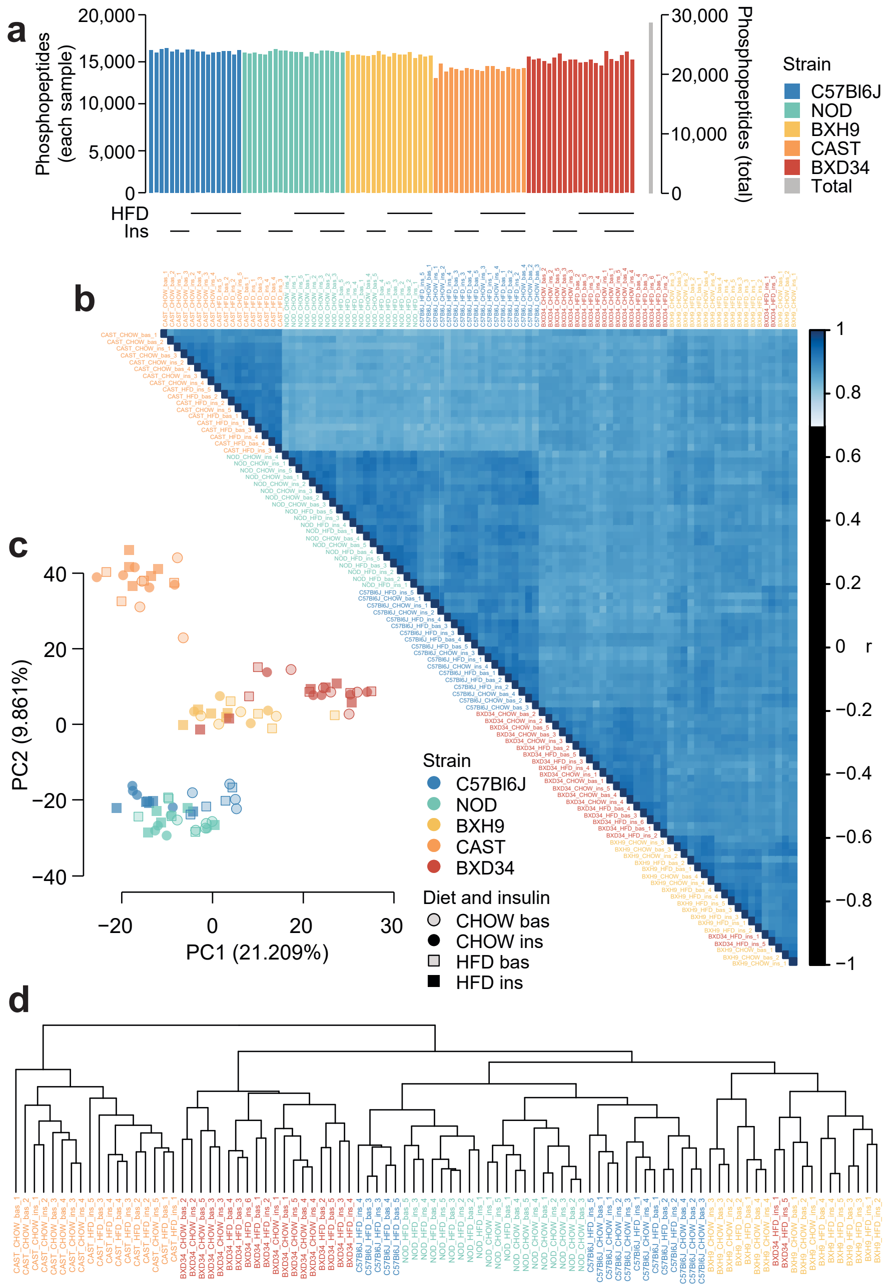


**Figure S2: Quality control analysis of phosphoproteomics data.** Related to Fig. 1. **a)** The number of unique class I phosphopeptides quantified in each sample and in total. **b)** Pearson's correlation was performed between each pair of samples. Samples are ordered by hierarchical clustering. **c)** Principal component analysis was performed on the phosphoproteome. The first two principal components (PC1 and PC2) are plotted for each sample and the percentage of overall variance explained by each principal component is indicated. "bas": unstimulated, "ins": insulin-stimulated. **d)** Hierarchical clustering was performed on all samples.

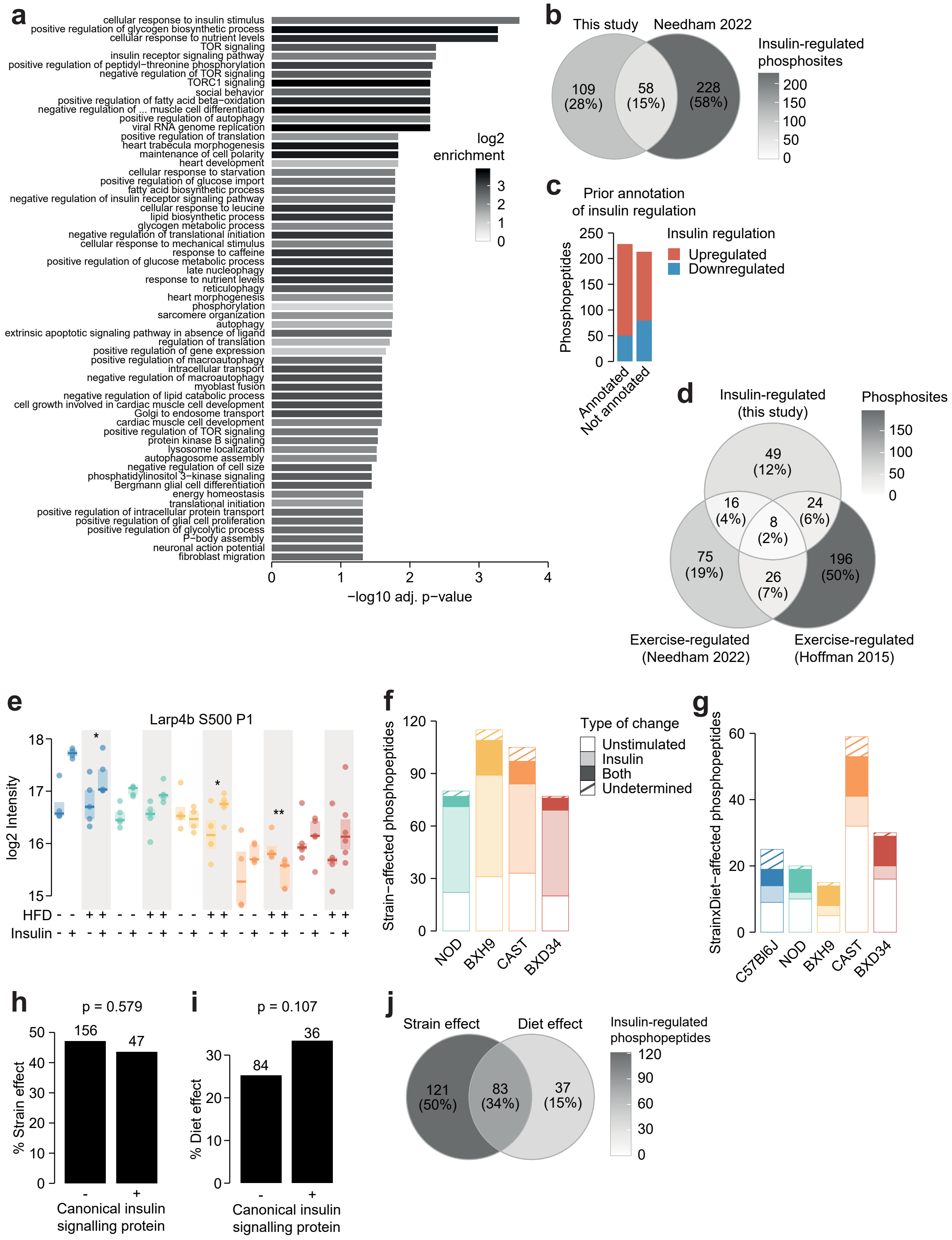


**Figure S3: Characterisation of the insulin-regulated phosphoproteome.** Related to Fig. 1-3. **a)** The enrichment of GO biological processes in genes containing insulin-regulated phosphopeptides relative to the entire phosphoproteome (one-sided Fisher's exact test, Benjamini-Hochberg p-value adjustment). Only significant pathways are shown (adj. $p < 0.05$). The pathway "negative regulation of vascular associated smooth muscle cell differentiation" is abbreviated. **b)** The number of phosphosites regulated by insulin in this study or a previous phosphoproteomic study of human skeletal muscle[1]. Only phosphosites quantified in both studies were considered. **c)** The number of insulin-regulated phosphopeptides with prior annotation of insulin regulation in the PhosphositePlus database[2]. **d)** The number of phosphosites regulated by insulin this study or regulated by exercise in two human phosphoproteomics studies[1,3]. Only phosphosites quantified in all three studies were considered. **e)** A phosphopeptide where HFD-feeding enhanced insulin responses in BXH9 but suppressed insulin responses in C57Bl6J and CAST. A two-way ANOVA was performed on insulin response values followed by two-sided t-tests comparing HFD to CHOW within each strain (q-values: *). **f)** Phosphopeptides with a Strain effect were examined to determine whether the effect was due to altered unstimulated phosphorylation ("Unstimulated"; Strain/C57Bl6J fold change > 1.3 in unstimulated samples), altered insulin-stimulated phosphorylation ("Insulin"; Strain/C57Bl6J fold change > 1.3 in insulin-stimulated samples), or both ("Both"). A proportion of phosphopeptides passed neither of these filters ("Undetermined"). **g)** The same analysis was performed on StrainxDiet-affected phosphopeptides, using the HFD/CHOW fold changes in either unstimulated or insulin-stimulated samples for each strain. **h-i)** The percentage of **h)** Strain effects and **i)** Diet effects (Uniform diet or StrainxDiet effect) among canonical or noncanonical insulin signalling proteins. P-values indicate two-sided Fisher's exact tests. The number of phosphopeptides in each group is shown. **j)** The overlap of Strain and Diet effects.

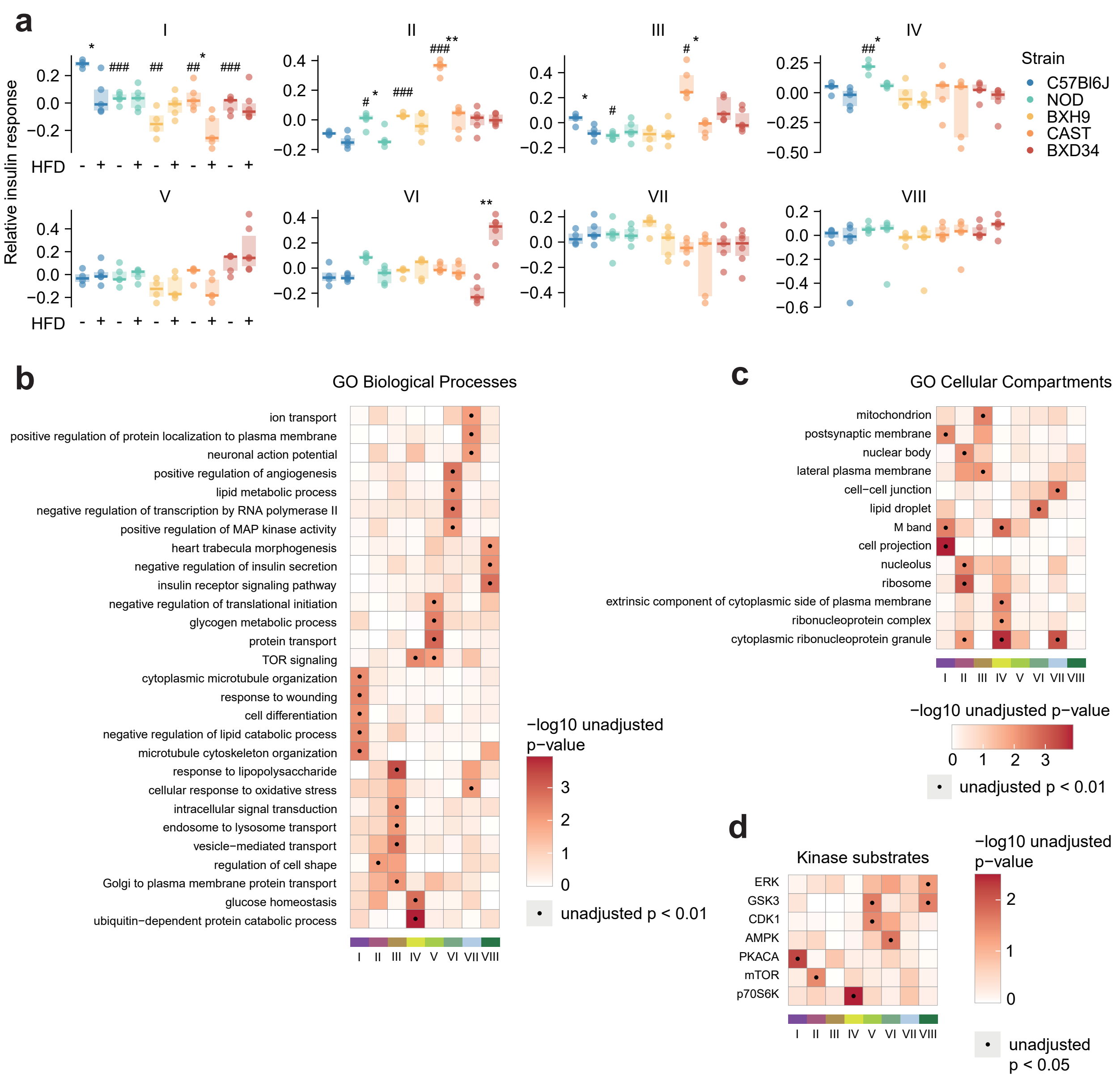


**Figure S4: Characterising insulin signalling subnetworks.** Related to Fig. 5. **a)** The eigenpeptides of each WGCNA-derived subnetwork. ANOVAs were performed on CHOW values following two-sided t-tests comparing each strain to C57Bl6J (Benjamini-Hochberg-adjusted p-values: #), and two-way ANOVAs were performed on all values followed by two-sided t-tests comparing HFD to CHOW within each strain (adjusted p-values: *). **b-d)** Rank-based enrichment of GO biological pathways, GO cellular compartments, and PhosphositePlus kinase substrates using phosphopeptide module membership scores with the "geneSetTest" function from the R package "limma". Membership score is defined as the absolute Pearson's correlation coefficient between a phosphopeptide's insulin response and the subnetwork's eigenpeptide. */#: $0.01 \leq p < 0.05$, **/##: $0.001 \leq p < 0.01$, ***/###: $p < 0.001$

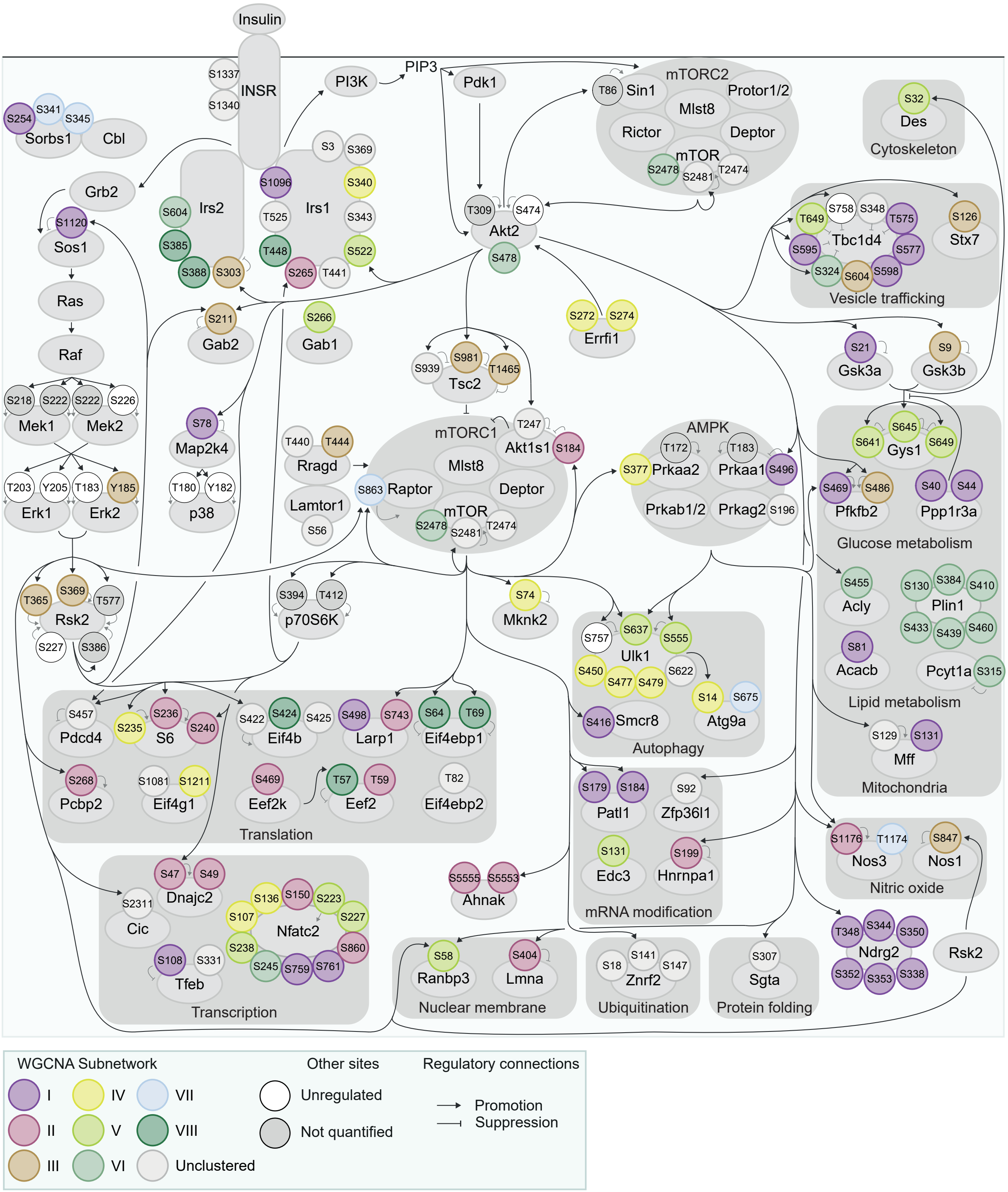


**Figure S5: Organisation of insulin signalling subnetworks.** Related to Fig. 5. The curated insulin signalling network displayed in Fig. 3 was annotated with WGCNA subnetworks from Fig. 5.

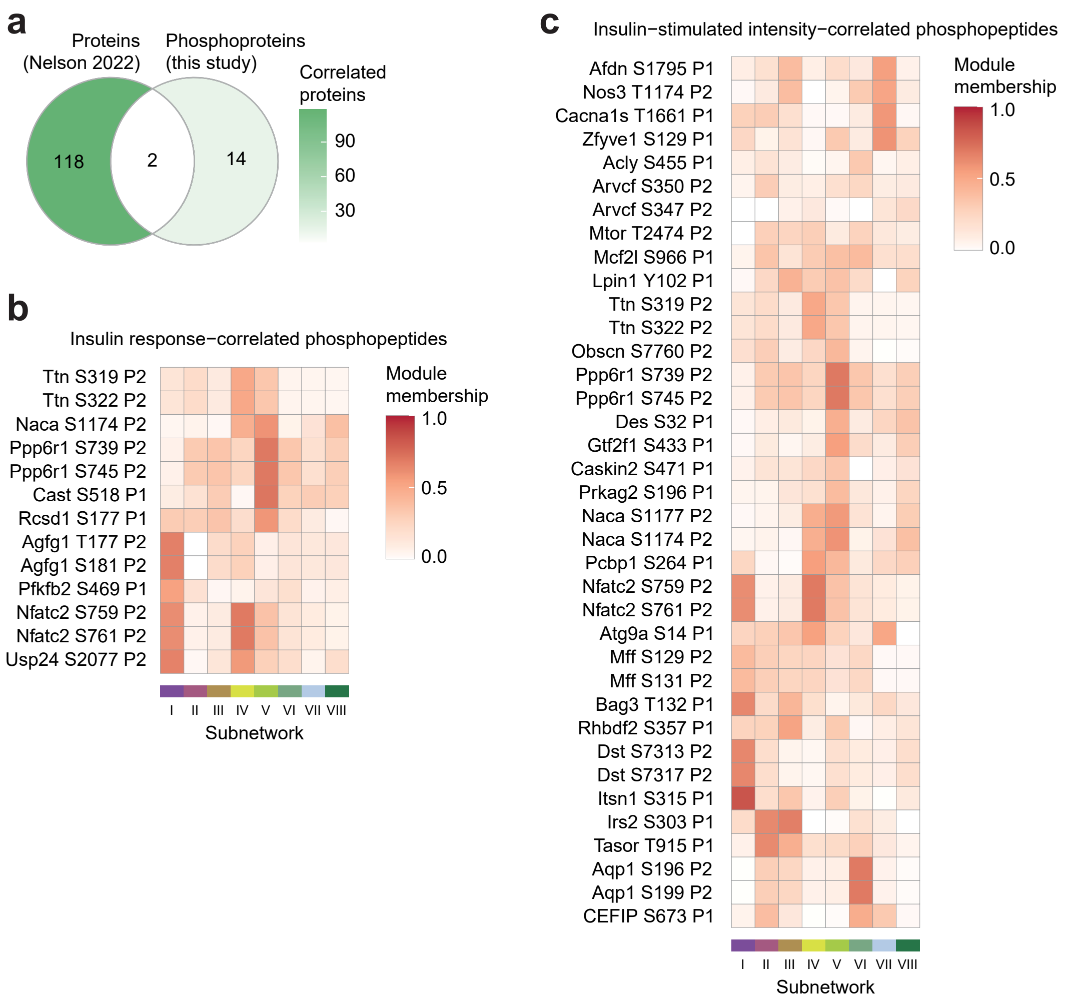


**Figure S6: Characterisation of glucose-uptake correlated phosphosites.** Related to Fig. 6. **a)** The intersection of proteins correlated with insulin-stimulated glucose uptake in the soleus of seven inbred mouse strains fed CHOW or HFD[4] ($p < 0.1$, $r > 0.35$ or $< -0.35$), with proteins containing glucose-uptake correlated phosphopeptides in this study. Only (phospho)proteins quantified in both studies are shown. **b-c)** The subnetwork membership scores for glucose-uptake correlated phosphopeptides using **b)** insulin-stimulated phosphopeptide intensity, or **c)** phosphopeptide insulin response values. Membership score is defined as the absolute Pearson's correlation coefficient between a phosphopeptide's insulin response and the subnetwork's eigenpeptide.

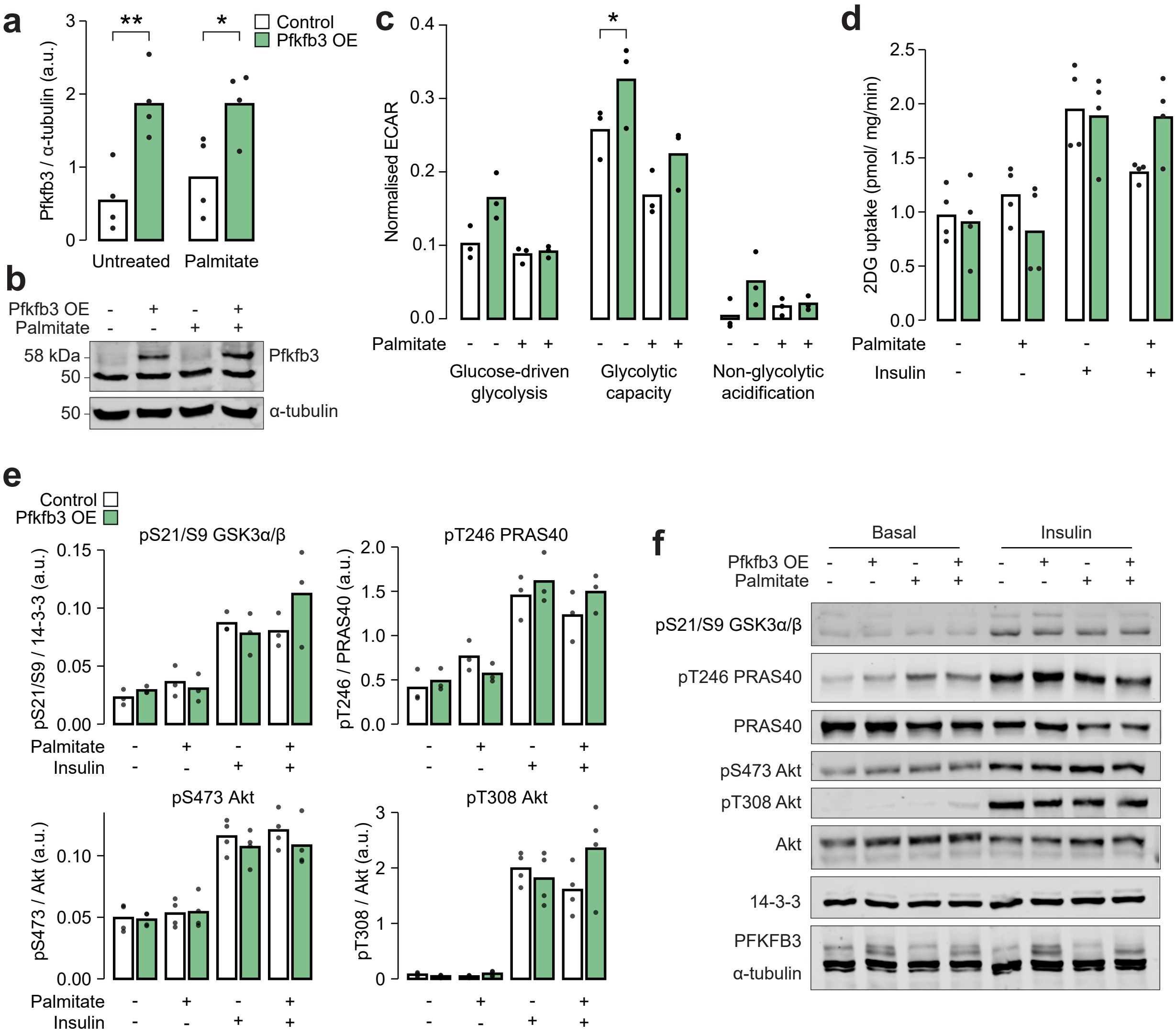


**Figure S7: Overexpression of Pfkfb3 enhances glycolytic capacity and reverses palmitate-induced insulin resistance.** Related to Fig. 6. **a)** Quantification and **b)** representative blot for immunoblotting of Pfkfb3 in L6-GLUT4-HA myotubes with or without Pfkfb3 overexpression, treated with palmitate (125 µM, 16 h) or BSA vehicle control. Two-way ANOVA was performed followed by Šidák's post-hoc tests assessing the effect of Pfkfb3 overexpression (*). n = 4 biological replicates. **c)** Extracellular acidification rate (ECAR) in L6-GLUT4-HA myotubes treated with glucose (10 mM, "Glucose-driven glycolysis"), oligomycin (5 µg/mL, "Glycolytic capacity"), or 2-deoxyglucose (50mM, "Non-glycolytic acidification"). A two-way ANOVA was performed followed by Tukey's posthoc tests comparing conditions within each of the three treatments (*). Not all significant comparisons are shown. n = 3 biological replicates. **d)** Unstimulated and insulin-stimulated glucose uptake (100 nM insulin, 20 min) in L6-GLUT4-HA myotubes. Insulin/unstimulated fold changes are shown in **Fig. 6h**. **e)** Quantification and **f)** representative blot for immunoblotting of insulin signalling phosphosites in unstimulated or insulin-stimulated (100 nM, 20 min) L6-GLUT4-HA myotubes.

*/#: 0.01 ≤ p < 0.05, **/##: 0.001 ≤ p < 0.01, ***/###: p < 0.001

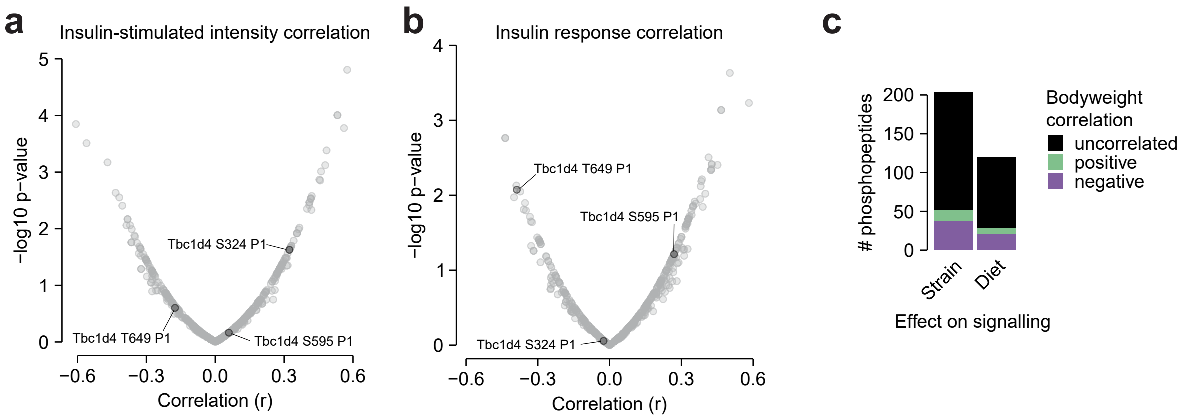


**Figure S8: Additional glucose uptake correlation analysis.** Related to Discussion. **a-b)** The correlation of insulin-stimulated glucose uptake with insulin-regulated phosphopeptides using **a)** insulin-stimulated phosphopeptide intensity, or **b)** phosphopeptide insulin response values, as displayed in **Fig. 6**. Canonical regulatory phosphosites on Tbc1d4 are indicated. The fourth canonical regulatory site S758 was not analysed due to insufficient quantification (quantified in 4/94 samples). **c)** The correlation of mouse bodyweight with the insulin responses of Strain or Diet-affected phosphopeptides either in CHOW-fed mice (Strain effects) or across both diets (Diet effects). The number of positively correlated (Pearson's correlation, $q < 0.1$, $r > 0.35$) and negatively correlated ($q < 0.1$, $r < -0.35$) phosphopeptides is shown.

***Table S1: Muscle phosphoproteomics***

(Page 1 "01_quantification") Normalized LFQ intensities of class I phosphopeptides. (Page 2 "02_analysis") Statistical analysis of phosphoproteome data.

***Table S2: Overlap between insulin and exercise-regulated phosphosites***

Insulin-regulated phosphosites from this study that were additionally regulated by exercise either in Needham et al.[1] or Hoffman et al.[3].

| *Phosphosite details* | | | | *Exercise regulation* | |
|---|---|---|---|---|---|
| **PhosphositePlus** | | | | | |
| **Site Group ID** | **Gene** | **Uniprot** | **Phosphosite** | **Hoffman 2015** | **Needham 2022** |
| 450254 | Eef2 | P58252 | T59 | + | + |
| 448040 | Eef2 | P58252 | T57 | + | + |
| 470754 | Mff | Q6PCP5 | S131 | + | + |
| 470753 | Mff | Q6PCP5 | S129 | + | + |
| 453632 | Larp1 | Q6ZQ58 | S498 | + | + |
| 455925 | Tbc1d4 | Q8BYJ6 | S324 | + | + |
| 4762876 | Svil | Q8K4L3 | S300 | + | + |
| 448238 | Gys1 | Q9Z1E4 | S645 | + | + |
| 480145 | Tns1 | E9Q0S6 | S1054 | + | |
| 3176152 | Ulk1 | O70405 | S450 | + | |
| 448064 | Rps6ka3 | P18654 | S369 | + | |
| 469782 | Dnajc2 | P54103 | S47 | + | |
| 486522 | Pcbp1 | P60335 | S264 | + | |
| 448094 | Rps6 | P62754 | S240 | + | |
| 448093 | Rps6 | P62754 | S236 | + | |
| 447594 | Mapk1 | P63085 | Y185 | + | |
| 453577 | Patl1 | Q3TC46 | S179 | + | |
| 447527 | Eif4ebp1 | Q60876 | S64 | + | |
| 482918 | Pcbp2 | Q61990 | S268 | + | |
| 4731870 | Speg | Q62407 | S542 | + | |
| 14570198 | Atg9a | Q68FE2 | S675 | + | |
| 455922 | Tbc1d4 | Q8BYJ6 | S348 | + | |
| 471364 | Lpin1 | Q91ZP3 | S923 | + | |
| 448239 | Gys1 | Q9Z1E4 | S649 | + | |
| 4693547 | Ttn | A2ASS6 | S322 | | + |
| 480195 | Usp24 | B1AY13 | S1296 | | + |
| 10782556 | Usp24 | B1AY13 | S1282 | | + |
| 483808 | Akap13 | E9Q394 | S2692 | | + |
| 3208338 | Akap1 | O08715 | S103 | | + |
| 3831732 | Tom1 | O88746 | S376 | | + |
| 2048085 | Rab12 | P35283 | S20 | | + |
| 447655 | Map2k4 | P47809 | S78 | | + |
| 4720237 | Lmod2 | Q3UHZ5 | T391 | | + |
| 450706 | Hspb6 | Q5EBG6 | S16 | | + |
| 4726016 | Tom1l2 | Q5SRX1 | S394 | | + |
| 4716961 | Slc20a2 | Q80UP8 | S316 | | + |
| 17544627 | Lmod1 | Q8BVA4 | S511 | | + |
| 11180047 | Tbc1d4 | Q8BYJ6 | S604 | | + |
| 455921 | Tbc1d4 | Q8BYJ6 | T649 | | + |
| 3195560 | Golga4 | Q91VW5 | S41 | | + |
| 485723 | Plekhf2 | Q91WB4 | S226 | | + |
| 4772204 | Ehbp1l1 | Q99MS7 | S1460 | | + |
| 470067 | Akt1s1 | Q9D1F4 | T247 | | + |
| 452430 | Plec | Q9QXS1 | S4620 | | + |

| | | | | | |
|---|---|---|---|---|---|
| 451009 | Ndrg2 | Q9QYG0 | T348 | | + |
| 468257 | Ndrg2 | Q9QYG0 | S338 | | + |
| 468259 | Ndrg2 | Q9QYG0 | S344 | | + |
| 447886 | Nos1 | Q9Z0J4 | S847 | | + |

***Table S3: Insulin signalling subnetworks***

WGCNA-derived subnetworks of insulin-regulated phosphopeptides.

***Table S4: Association of kinase enrichment with insulin-stimulated glucose uptake***

Pearson's correlation of KSEA enrichment scores with insulin-stimulated glucose uptake for all kinases with Strain or Diet effects. Correlation was performed on all values or on the medians of each Strain-Diet combination.

| | ***Correlating all values*** | | ***Correlating Strain-Diet medians*** | |
|---|---|---|---|---|
| **Kinase** | **r** | **p** | **r** | **p** |
| SGK | 0.026282557 | 0.85773407 | -0.114513813 | 0.752760165 |
| Aur | 0.181856398 | 0.221176679 | 0.446186208 | 0.196168185 |
| p90RSK | 0.042637179 | 0.771137201 | 0.392619919 | 0.261741297 |
| aPKC | 0.020857899 | 0.886881337 | 0.059269839 | 0.870801728 |
| GSK3 | -0.023496391 | 0.872683503 | 0.200939915 | 0.577765996 |
| CDK5 | 0.092355979 | 0.527940053 | 0.136553891 | 0.706796386 |
| P38 | 0.186315084 | 0.199914602 | 0.202333014 | 0.575075383 |
| CDK1 | 0.134654975 | 0.356287427 | 0.245970422 | 0.493328434 |
| CK2 | 0.062152944 | 0.671383691 | -0.206449396 | 0.567152858 |
